# The Reuse-and-Append Memory Principle: Application to Latent Cause Inference

**DOI:** 10.64898/2026.09.09.750332

**Authors:** Zied Ben Houidi, Samuel J. Gershman

## Abstract

The latent cause theory of memory modification provides a computational account of how the brain decides whether to update existing memories or form new ones, but leaves unspecified the neural mechanisms implementing this inference. We propose Reuse-and-Append Memory (RAM), a set of mechanistic principles that achieve the same goal through sparse neural coding, Hebbian learning, and pattern-matching dynamics. The core idea is that neurons encoding prior experiences are automatically reactivated by similar stimuli, while uncommitted neurons are recruited to encode genuinely novel aspects of each experience, including the passage of time. We present a computational model instantiating these principles and show that it reproduces acquisition, extinction, renewal, and spontaneous recovery in fear conditioning. By committing to a neural mechanism, we found that RAM separates what is typically modeled as a single prediction error driving memory updating or formation into two independent signals: an immediate novelty signal that emerges from coverage-based allocation of new neurons, and an outcome mismatch signal that later activates safety circuits when an expected outcome fails to arrive. RAM also reproduces the dependence of recovery on the timing of stimulus reminders (the Monfils-Schiller effect), but predicts it to be inherently fragile, consistent with the mixed empirical record. RAM suggests that the Monfils-Schiller effect, often attributed to reconsolidation, leaves the original fear memory intact: a bridging event carries safety information to new contexts through offline co-retrieval. More broadly, none of the phenomena we explain require modifying existing memories, and the most stable configuration is the extreme form of the principle: always append, never overwrite.

---

> No man ever steps in the same river twice
>
> Heraclitus

## 1 Introduction

The latent cause theory of memory modification (Gershman et al., 2017) offers a computational account of how memories change through experience. According to this framework, the brain performs Bayesian inference to determine whether new sensory observations arise from the same latent cause as previous memories, triggering in-place modification, or from a new cause entirely, triggering formation of a separate memory trace. This explains why fear memories return after extinction (spontaneous recovery), why extinction in a new context fails to generalize (renewal), and why a single retrieval trial before extinction can prevent fear recovery (Monfils et al., 2009; Schiller et al., 2010). Yet the theory does not specify how biological circuits carry out this inference. What neural machinery segments experience into distinct causal clusters? How do neurons approximate inference over latent causes given the constraints of sparse firing and local synaptic plasticity?

We propose Reuse-and-Append Memory (RAM) as a set of mechanistic principles that achieve the same goal. When a new experience shares features with a prior one, the neurons that encoded those features are activated again (*reuse*). When features are genuinely novel, including changes in context or the passage of time, uncommitted neurons (e.g., those freed by synaptic decay of unrehearsed memories) are recruited to encode them (*append*). This creates a hypothesized architectural constraint: *if* each experience includes novel episodic components requiring new neurons, and *if* these new recruits are more plastic than established ones, then the system cannot instantaneously overwrite existing patterns.

Consider a tone presented today versus yesterday: while neurons encoding the tone itself may overlap, the complete neural pattern necessarily differs due to distinct time cells, changing internal states, and unique contextual elements. No two experiences, however similar, produce identical sparse hippocampal representations; we never swim in the same neural river twice. This division of labor (reused neurons maintain stability, new neurons enable learning) ensures that each episodic memory forms through Hebbian binding of sparse neurons co-active in the same moment. At retrieval, which sparse pattern is activated determines what memory and its associated outcome are expressed.

Consider extinction, where the conditioned stimulus (CS; e.g., a tone) is presented without the unconditioned stimulus (US; e.g., a shock). In latent cause theory, extinction creates a new latent cause because large prediction errors signal a different CS-US joint distribution. In RAM, extinction creates a new event because the extinction experience (same CS, different time, different context, no US) cannot fully reuse the acquisition ensemble. Coverage is low because the cue contains too many features that do not match the acquisition memory. A new event is allocated automatically, new neurons bind with a safety outcome, and the original fear memory remains intact.

This principle extends to the other phenomena that motivated latent cause theory. Renewal emerges from pattern completion: when retrieval cues match the acquisition ensemble, that memory activates. Spontaneous recovery reflects the transience of episodic memories: synaptic turnover gradually weakens unrehearsed traces, a process that also frees neurons for new encoding. Extinction memories fade over time, while fear is additionally maintained by a direct CS-to-US pathway not subject to the same turnover. Simulations reveal that the sensitivity of recovery to the timing of a CS reminder (the Monfils-Schiller effect, described further below), often attributed to reconsolidation, can occur without modifying the original fear memory: a bridging event carries safety information to new contexts through offline co-retrieval. The model predicts this effect to be inherently fragile, dependent on intensive offline replay, which may explain why individually housed animals show stronger effects than group-housed ones (Kredlow et al., 2016).

In this paper, we formulate the RAM principles, ground each component in neural evidence, and establish a structural correspondence with latent cause theory (Section 2). We present a computational model (Section 2.2) that robustly reproduces these phenomena across wide parameter ranges (Section 4). By committing to a circuit-level abstraction, we reveal structure previously not visible at the computational level: what is typically modeled as a single prediction error separates into two independent signals, a novelty signal from coverage-based allocation that creates the memory, and an outcome mismatch signal that activates safety circuits when an expected outcome fails to arrive. We discuss other predictions and connections to related work in Section 5.

## 2 The Reuse and Append Memory Principle

RAM is a principle about how biological memory encodes, preserves, and retrieves past experiences. In this paper, we develop it in the setting of Pavlovian fear conditioning. This section presents the conceptual framework (Section 2.1), provides the computational model (Section 2.2), establishes a structural correspondence with latent cause theory (Section 2.3), and finally reviews the biological grounding for each component (Section 2.4).

### 2.1 RAM Overview

Experiences are encoded as sparse neural patterns. When a new experience shares features with a prior one, the neurons that encoded those features (e.g. tone-selective neurons in auditory cortex, context-selective neurons in hippocampus) are activated again. This *reuse* is automatic, a consequence of neural tuning rather than a decision. When features are genuinely novel (time has progressed, a new context or stimulus has appeared), uncommitted neurons are recruited to encode them. We call this *append*. The resulting pattern partially overlaps with prior patterns; the shared neurons reflect what is familiar while the newly recruited neurons reflect what is new.

Since synaptic modification is activity-dependent and representations are sparse, the vast majority of synapses belonging to prior memories are inactive. Existing memories are therefore not overwritten. Furthermore, over time, consolidation stabilizes new traces and reduces their plasticity, while unconsolidated traces may decay, potentially creating a new pool of neurons to encode novel representations. Co-active feature neurons are bound together by higher-order *event neurons* that encode a specific group of features as a coherent episode. Multiple events can share feature neurons through reuse while remaining distinct, because each is bound by a different event neuron. The result is a hierarchy: feature neurons encode individual stimuli and contexts, while event neurons encode the particular combinations in which they co-occurred.

When the animal later encounters a similar cue, stored event representations compete for activation based on how well each matches the incoming input, and the winning event drives behavior.

#### 2.1.1 Two reference frames for partial overlap

The retrieval competition just described requires running acue against stored event representations. This interaction between the cue and existing events will answer two distinct questions.

The first is *retrieval*: how strongly is each stored event engaged by the current input? This is measured by overlap, an event-centric computation: what fraction of the stored event’s feature neurons are active in the cue. An event whose representation involves ten feature neurons, eight of which are active in the cue, is strongly retrieved regardless of how many additional neurons the cue contains.

The second question is *allocation*: is the current experience adequately captured by an existing event representation, or does it require a new one? This is measured by coverage, a cue-centric computation: what fraction of the cue’s active feature neurons belong to an existing event. A cue with twenty active feature neurons, eight of which match a stored event, has only 40% coverage; substantial novelty remains “unexplained”. Hence, low coverage triggers the recruitment of a new event neuron to bind the current group of features into a new event representation.

The asymmetry between these reference frames matters. If allocation used the event-centric frame, any fully reactivated event would appear to explain the cue regardless of what else it contained, and the animal would never form new memories for experiences richer than what it has already stored. If retrieval used the cue-centric frame, a rich stored event would fail to be retrieved whenever the cue contained only a fraction of its features, and pattern completion from partial cues would break down. Figure 1 illustrates both frames. As a computational convenience, not a literal mechanism, Section 2.2 formalizes them as set-intersection measures over sparse neural pools, differing only in the denominator.

**Figure 1:**
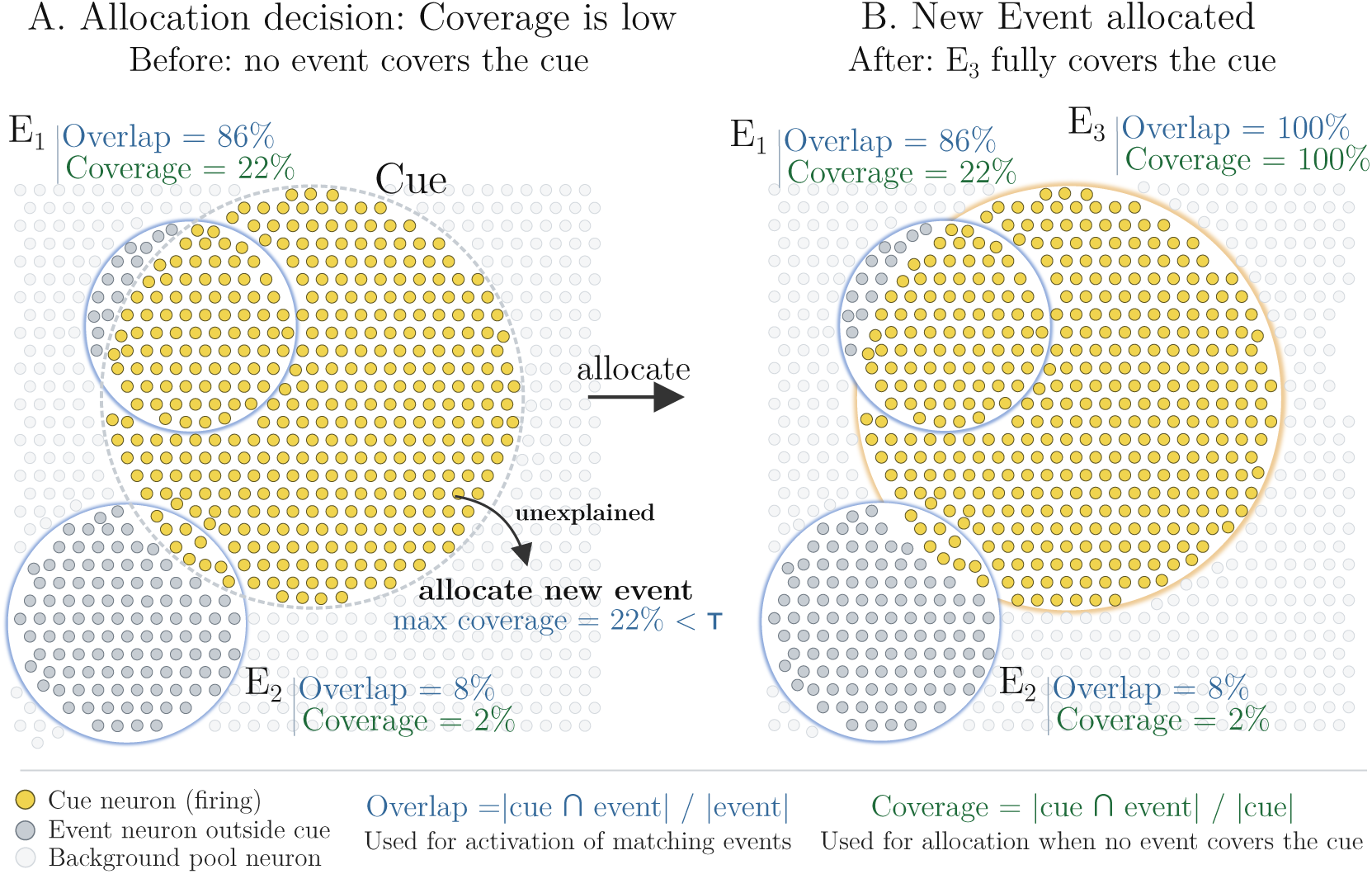
Illustration of the coverage and overlap metrics. A new Cue has a high overlap with a previous Event *E*_1_ but not *E*_2_. *E*_1_ is hence retrieved more strongly. None of the existing events has a high coverage with the current cue, so a new Event *E*_3_ is created.

#### 2.1.2 Illustration through fear conditioning

Figure 2 illustrates these principles through a canonical protocol.

**Figure 2:**
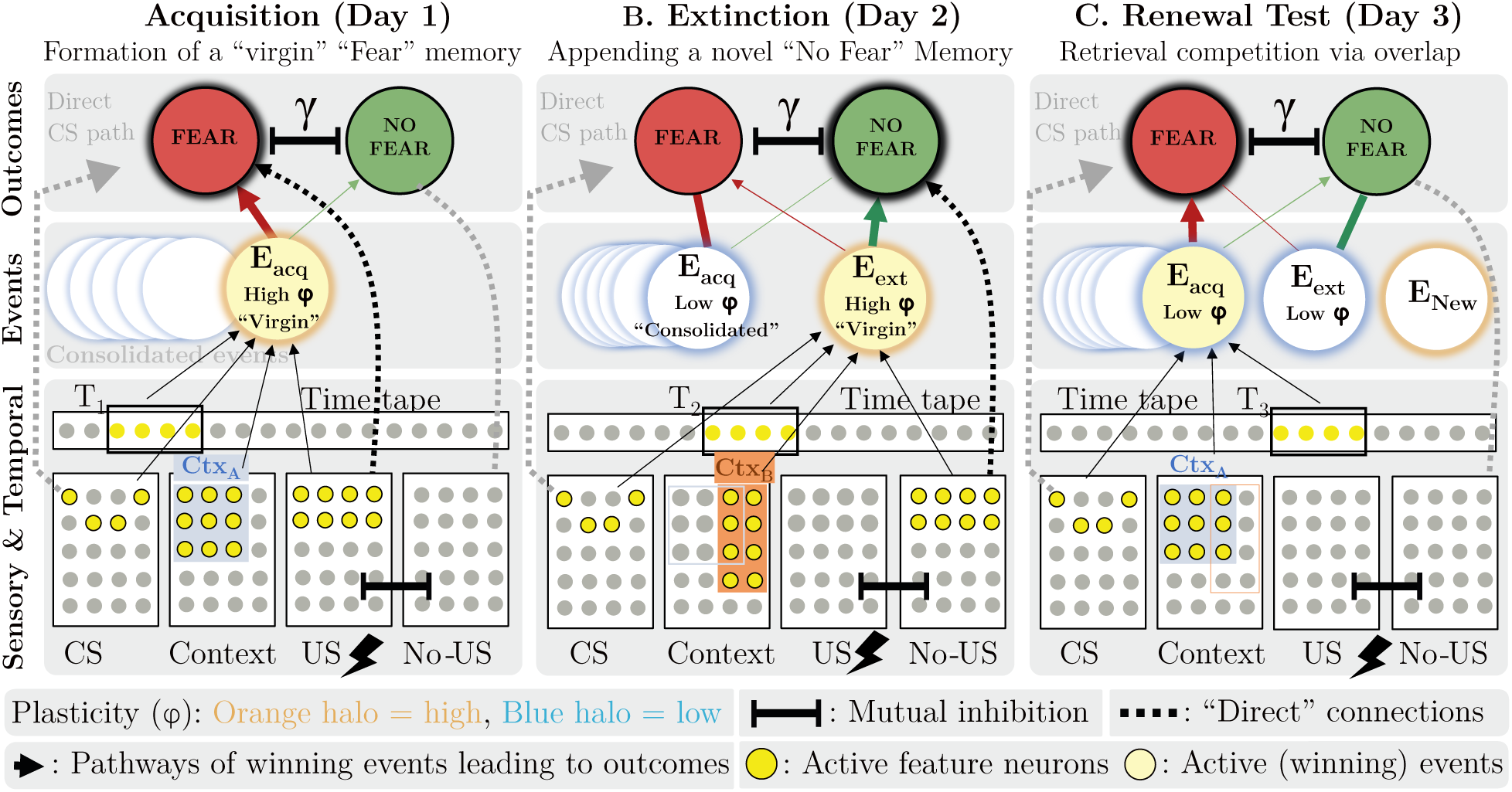
Overview of RAM dynamics across acquisition, extinction, and renewal. **Panel A (Acquisition)**: A tone (CS) paired with shock (US) in Context A creates event *E*_acq_. Active features bind together; co-occurrence with shock drives *E*_acq_ → Fear learning. New events have high plasticity (*ϕ*, orange). **Panel B (Extinction)**: The CS without shock in Context B creates a new event *E*_ext_, because time and context have changed and existing memories inadequately capture this experience. *E*_acq_, now consolidated (blue, low *ϕ*), remains intact. Absence of expected shock activates No-US features, driving *E*_ext_ → No-Fear learning. The original fear memory is not over-written. **Panel C (Renewal):** The CS in Context A overlaps more with *E*_acq_ (matching CS and context) than with *E*_ext_ (matching CS only). E_acq_ wins retrieval competition, driving fear expression despite prior extinction.

During **acquisition** (Figure 2A), a tone (CS) paired with shock (US) in Context A co-activates neurons from distinct feature pools: tone-selective, context-A, temporal, and US-responsive neurons. Coverage by existing memories is low (the combination is novel), so a new event representation, *E*_acq_, is allocated. Hebbian co-activation with shock drives a strong *E*_acq_ → Fear association. Overnight consolidation stabilizes these synapses and reduces their plasticity.

During **extinction** (Figure 2B), the CS is presented without shock in Context B. The tone neurons fire again (reuse), but context has changed, time has advanced, and no shock arrives. Coverage by *E*_acq_ is low: too many features are unexplained. A new event, *E*_ext_, is allocated. Because *E*_acq_ is partially retrieved, pattern completion reactivates its US neurons top-down, generating an expectation of shock. When no shock arrives, the mismatch between expected and actual outcome activates a safety signal through mutual inhibition between US-responsive and US-absence-detecting neural populations (Section 2.2.9). These safety neurons are incorporated into *E*_ext_ and drive a No-Fear association. *E*_acq_’s synapses are unchanged: its partners from acquisition are mostly inactive, and its consolidated connections resist modification.

At **renewal** (Figure 2C), the CS is presented in Context A. *E*_acq_’s overlap with the cue is high (both CS and context-A neurons match); *E*_ext_’s overlap is low (only CS neurons match). *E*_acq_ dominates retrieval and drives Fear. A direct CS-to-Fear pathway, consolidated during acquisition and resistant to modification (Doyère et al., 2003), additionally reinforces this outcome. Fear returns not because extinction was undone (*E*_ext_ and its safety association remain intact) but because the balance of retrieval shifted.

#### 2.1.3 Weight decay and inter-event linking through offline replay

The remaining phenomena (spontaneous recovery and the Monfils-Schiller timing effect) involve additional dynamics: episodic weight decay and inter-event linking through offline replay.

Weight decay models the progressive weakening of synaptic connections in memories that are not rehearsed or consolidated, causing extinction memories to fade while consolidated fear memories persist. Inter-event linking occurs during offline silence periods, where the most salient recent memories (events) are replayed in sequence. Event memories that fire together wire together, thus creating, if consolidated, durable associative links. These phenomena are formalized in Section 2.2 and discussed in Section 4.

### 2.2 Computational Model

This section presents one concrete instantiation of the RAM principle, framing each component as an approximation that captures the essential dynamics while remaining computationally tractable.

#### 2.2.1 Representations

Features are implemented as disjoint random sparse pools. Each feature family occupies a separate subset of neurons; instances within a family activate exactly k neurons from an available pool of n. This ensures that overlap between events arises only from shared semantic content.

The feature families are: CS (*n*_cs_, *k*_cs_), context (*n*_ctx_, *k*_ctx_), time (*n*_time_, *k*_time_), US (*n*_us_, *k*_us_), No-US (*n*_nous_, *k*_nous_), and silence (*n*_sil_, *k*_sil_). The No-US features encode the active detection of US absence (Section 2.2.9). Silence features activate during inter-trial periods when no external stimulus is present.

Time is represented via a non-wrapping tape: at step *t*, active time neurons span [*t, t* + *k*_time_). Experiences separated by more than *k*_time_ steps share no temporal features. Context similarity between environments is controlled by an overlap parameter *ρ*_ctx_∈ [0, 1] specifying the fraction of shared context neurons.

#### 2.2.2 Retrieval and Allocation Metrics

The retrieval and allocation computations from Section 2.1 are implemented as set-intersection measures over sparse neural patterns, differing only in normalization (as illustrated in Figure 1).

**Overlap** (memory-centric, for retrieval):

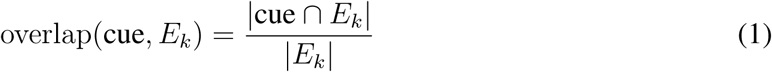

**Coverage** (cue-centric, for allocation):

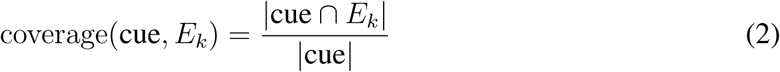

**Retrieval competition**. Events compete for activation via softmax:

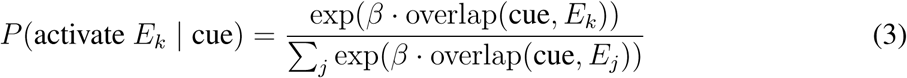

Softmax is adopted as one concrete monotonically increasing competition function, with temperature parameter *β* controlling sharpness. Any function that increases with overlap would serve the same role. Section 4.4 shows the model is insensitive to *β* across the range {1, . . ., 12} under the retained paper parameters.

**Allocation decision.** A new event is allocated when the best-matching existing memory has insufficiently high coverage:

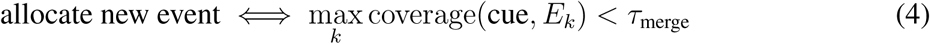

**Overlap weighting.** In the simulations, each feature family (CS, context, time, US, No-US) contributes to the overlap and coverage computations with its own weight w_f_, replacing set cardinalities with weighted sums (Appendix B). A low w_time_ reduces the influence of temporal mismatch, allowing memories formed at different times to compete primarily on “semantic” content.

**Interpretation.** We present coverage and overlap as set-intersection measures because this makes the model tractable, but we do not claim the brain computes either quantity explicitly. Instead, they could emerge from network dynamics: in a sparse competitive network with Hebbian-trained weights, familiar input patterns preferentially activate the ensembles whose weights are tuned to them, and input current flows into those pre-existing networks. Input components that match no established ensemble would leave residual activity that falls on whichever neurons are currently most excitable. We believe such residual activity is not separately computed, but is what remains once familiar ensembles have activated their share, making coverage and overlap two views of the same phenomenon. The parameters τ_merge_, β, and the family weights w_f_ should be thus read as effective parameters of this approximation rather than quantities the circuit evaluates.

#### 2.2.3 Outcome Competition

Fear and No-Fear outcomes are modeled as mutually inhibiting populations inspired by Wilson-Cowan dynamics (Wilson and Cowan, 1972). Starting from *F* = *N* = 0, the following update equations are iterated for T steps (*T* = 6):

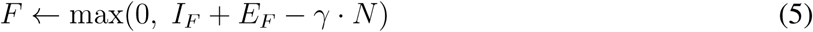

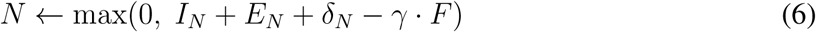

where 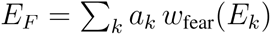 and 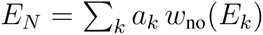 are summed inputs from retrieved events (excluding newborn events, which carry no prior learning), weighted by their fear and safety associations, and γ controls mutual inhibition strength. The intrinsic drives are

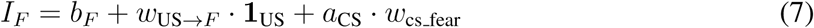

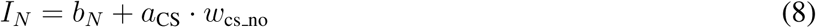

where *b_F_* and *b_N_* are tonic biases (*b_N_* provides a baseline safety signal), *w*_US→_*_F_* is the innate US-to-Fear drive (active when the US is present), and a_CS_ is the overlap between the cue and the stored CS pool, which activates the CS pathway (Section 2.2.6). The No-US bump δ*_N_* (Section 2.2.9) is applied as a separate input to the No-Fear channel when the safety vote dominates.

#### 2.2.4 Plasticity and Consolidation Under Novelty Gating

Each event maintains a plasticity coefficient *ϕ* ∈ [*ϕ*_min_, 1] that modulates learning rate. Newly allocated events start with *ϕ* = 1 (high plasticity); repeated activation causes *ϕ* to decay toward *ϕ*_min_. Learning uses two rates: *η*_ext_ = 0.3 for external trials (CS orUS present) and *η*_imag_ = 0.08 for silence periods (offline rehearsal). All active events learn proportionally to their activation level:

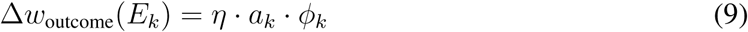

where *a_k_* is the event’s activation and η ∈ {*η*_ext_, *η*_imag_} depending on whether external input is present. Weights are clipped at symmetric caps (*w*_fear max_ = *w*_no max_).

Events accumulate a consolidation budget proportional to total learning. When this budget exceeds a threshold, the event becomes *permanent*: its *ϕ* is frozen at *ϕ*_min_, protecting it from further modification. Conversely, events that fail to consolidate within a critical window (*T*_virgin_ = 20 steps) are pruned. This two-level system (continuous *ϕ* modulation plus discrete consolidation) captures both short-term metaplasticity and long-term memory stabilization.

#### 2.2.5 Lability and Reconsolidation

The model includes an optional reconsolidation mechanism: when a consolidated event’s activation exceeds a lability threshold, its plasticity is temporarily re-opened (*ϕ* → 1), allowing modification. We disable this mechanism by default since our simulations show that the five target phenomena emerge without reconsolidation. Furthermore, Section 4.4 shows that enabling lability introduces a dependence on retrieval sharpness (*β*) that the baseline model avoids. At the paper’s baseline *β* = 1, lability permanently corrupts the fear memory during extinction.

#### 2.2.6 CS Pathway

In addition to event-level associations, the model includes a direct CS → Fear / CS → No-Fear pathway that bypasses episodic event representations. This pathway operates independently of event memories: it maintains its own weights (*w*_cs fear_, *w*_cs no_), plasticity state, and consolidation dynamics. Both weights are symmetrically capped at *w*_cs fear max_. When a CS is present in a cue, the overlap *a*_CS_ between the cue and the stored CS pool activates the pathway, driving Fear and No-Fear in proportion to *w*_cs fear_ and *w*_cs no_ (Section 2.2.3). The biological motivation for this component, a direct thalamo-amygdala projection, is reviewed in Section 2.4.

During acquisition, the CS pathway learns a strong CS → Fear association with full plasticity. After overnight consolidation, plasticity drops to zero (*ϕ*_cs min_ = 0) and the pathway is frozen. This is an idealization: the framework accommodates any degree of persistence; but we explore the frozen endpoint because it provides the most robust spontaneous recovery (Section 4.2). In the end, the key requirement for spontaneous recovery is that *some degree* of CS pathway persistence outlasts episodic memory decay.

#### 2.2.7 Weight Decay

Consolidated event weights decay slowly during silence periods, modeling progressive synaptic weakening in unrehearsed memories:

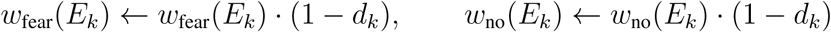

where *d_k_* is the per-step decay rate. US-bearing events (those containing shock neurons) decay at a separate rate *d*_us_. Non-US events decay at rate *d_k_* = *w*_decay rate_.

Spontaneous recovery requires that fear outlasts extinction over long delays. Two independent mechanisms can produce this asymmetry: the frozen CS pathway (Section 2.2.6), and slower decay of traumatic (US-bearing) memories relative to extinction memories. Either mechanism alone is sufficient. In our baseline configuration, we set *d*_us_ = 0, but the model successfully passes all tested phenomena for *d*_us_ up to 0.003 (Section 4.5). The result is a hierarchy where episodic extinction memories gradually weaken while fear persists through the CS pathway and the acquisition event’s slower decay.

#### 2.2.8 Silence, Rehearsal, and Links

##### Offline replay

During silence periods (no external input), the model rehearses recent events. A pool of events is selected based on recency-weighted activation traces (tags), and these events are reactivated sequentially. Each rehearsed event is partially reactivated: only a quarter of its stored neurons are sampled into the cue, modeling the degraded nature of offline replay. This partial cue nevertheless triggers pattern completion and retrieval, and the retrieved events enter the same outcome competition and weight-update pipeline as external trials, at a reduced learning rate (*η*_imag_; Section 2.2.4). A fear event that is replayed thus reactivates its shock associations and reinforces its fear weight; an extinction event reinforces its safety weight. Rehearsal is limited to *T*_replay_ consecutive steps per silence period; beyond this cap, silence is quiet and only weight decay operates.

##### Inter-event links

Two Hebbian linking rules operate during offline rehearsal. Co-activity links strengthen when two events are simultaneously active above a threshold:

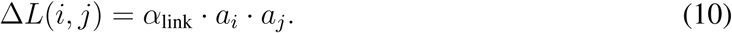

Sequential links strengthen between events rehearsed on consecutive silence steps:

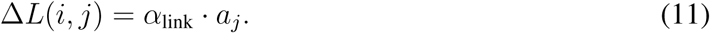

Both are bidirectional and form *only* during silence, never during external trials. Section 2.4 reviews evidence that inter-memory co-reactivation occurs preferentially during quiet wakefulness (Zaki et al., 2025) and that extinction trials actively suppress fear engram reactivation (Lacagnina et al., 2019). These links provide a retrieval bonus: when one event is activated, its linked events receive additional activation. Our simulations show that this mechanism is necessary for the Monfils-Schiller effect (Section 4.3), where bridge events formed during retrieval become linked to the acquisition memory through offline replay.

#### 2.2.9 No-US Detection: Safety Signal Through Expectation Violation

Safety learning requires more than the passive absence of shock; shock is absent during every non-shock moment of life. What the model needs is an active safety signal that fires specifically when shock was *expected* and did not arrive. In our framework, pattern completion provides the expectation: when a fear event is retrieved, its stored US features are reactivated, generating a prediction of shock. If no shock arrives, this unconfirmed expectation triggers the creation of No-US features in subsequent events.

Concretely, when events are retrieved, those encoded with US neurons contribute to a US vote, and those encoded with No-US neurons contribute a competing safety vote. When the US vote dominates but no shock arrives from external input, the violated expectation triggers No-US feature creation in subsequent events. Whenever the safety vote dominates, the No-US bump δ_N_ is applied to the No-Fear channel of the outcome competition (Section 2.2.3). Section 2.4 reviews a possible biological implementation of this mechanism through mutually inhibiting neural populations.

Two properties emerge from this mechanism without additional rules. First, safety learning is delayed at the onset of extinction. At the first CS-alone trial in a new context, the acquisition event is partially retrieved and establishes a US prediction. But this expectation was not yet present when the trial’s cue event was constructed, so the first extinction event is born without No-US markers. From the second trial onward, the established expectation is violated by the absent US, and new events incorporate active No-US features. Second, memories whose associations have fully decayed contribute nothing to the vote. An event with near-zero weights carries no predictive content, generates no expectation, and triggers no surprise at the absence of shock.

#### 2.2.10 Processing Pipeline

Algorithm 1 summarizes how the components above compose into a single processing step.

### 2.3 Correspondence with Latent Cause Inference

Latent cause theory (Gershman et al., 2017) rests on two principles: simplicity (observations tend to arise from few causes) and contiguity (temporally close events likely share a cause). The brain assigns each experience to a latent cause through Bayesian inference, creating a new cause when no existing one is sufficient. RAM instantiates both principles through sparse pattern dynamics without explicit probability computations. Preferential reuse of existing neurons provides the simplicity bias: a new event is allocated only when existing memories inadequately cover the cue. Temporal feature overlap provides contiguity: events close in time share active neurons, making them more likely to activate the same memory. Table 1 summarizes the correspondence.

**Table 1:**
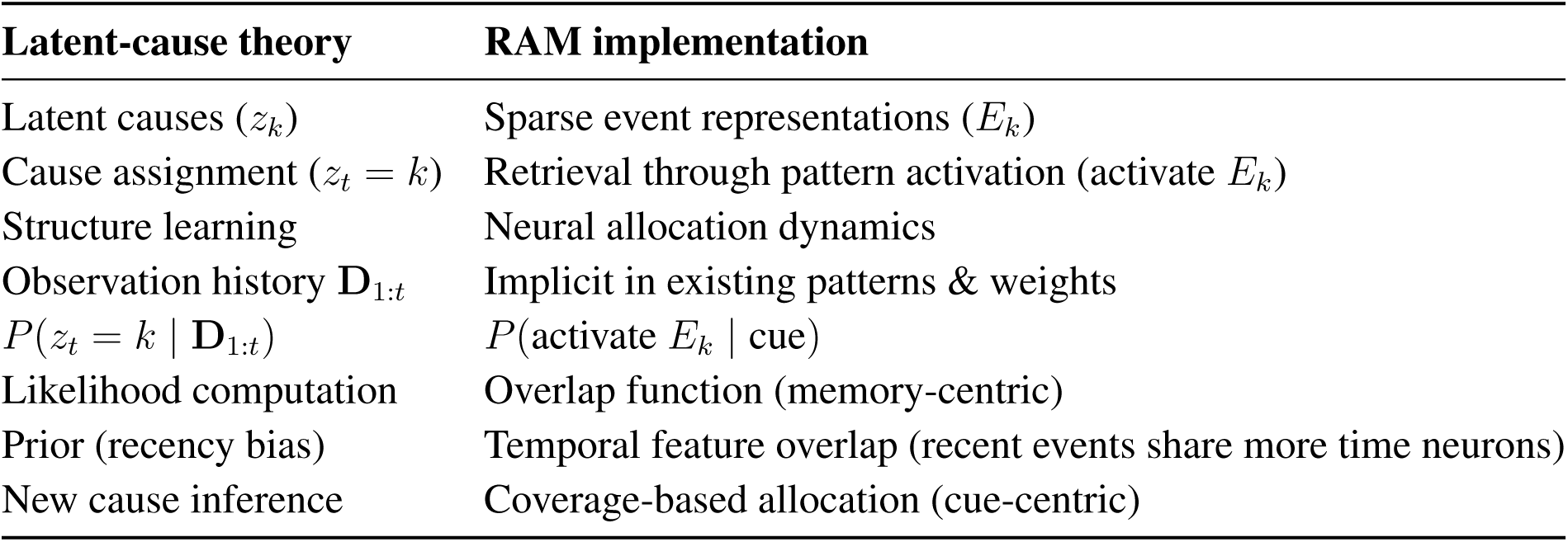
Mapping latent-cause theory to Reuse-and-Append Memory.

In Section 2.2, stored events compete for activation via a monotonically increasing function of overlap. When this function is instantiated as softmax (Eq. 3), the overlap function parallels the role of the log-likelihood in Bayesian cause inference: it measures how well an existing memory accounts for the current observation. The coverage function, used for allocation, plays the role of assessing whether any existing cause adequately explains the data. Note that this mapping is structural rather than exact, since continuous posteriors and discrete sparse patterns are different mathematical objects. Here, softmax is one possible implementation, chosen for convenience, but the model’s insensitivity, under the retained paper parameters, to β across the range {1, . . ., 12} (Section 4.5) suggests that the phenomena depend more on overlap-based values themselves.

#### Algorithm 1

One processing step of the RAM model.

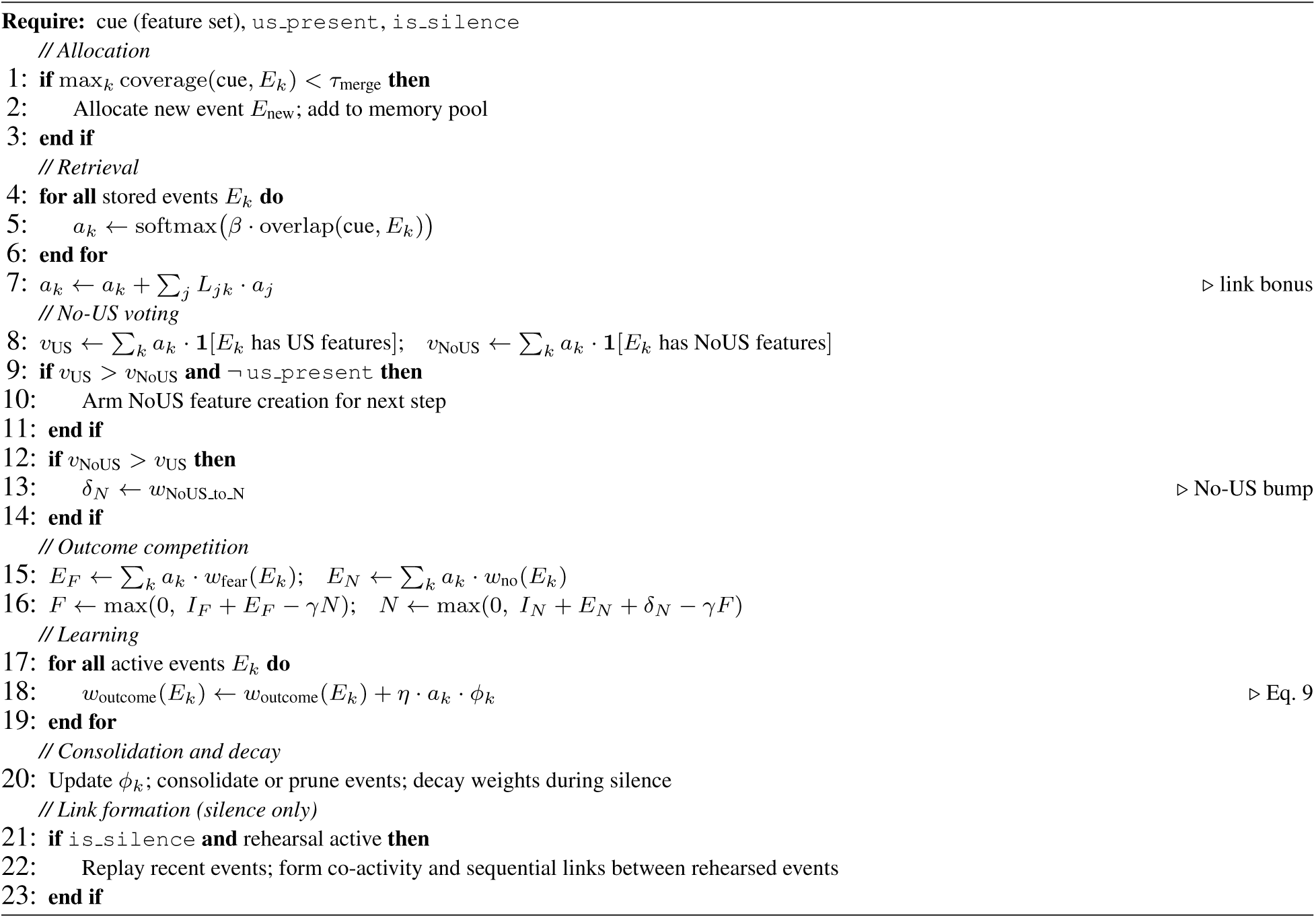

#### Two signals in place of prediction error

Latent cause theory assigns prediction error a dual role: signaling that the situation is novel and that an expected outcome did not occur. Committing to the RAM principle decomposes this into two independent signals.

The first is a *novelty signal* arising from allocation. When existing memories poorly cover the current cue, new neurons are “automatically” recruited. The proportion of newly recruited neurons in the resulting pattern is hence somewhat immediately available as a local readout of novelty.

The second is an *outcome prediction error* arising from violated expectations. When retrieved events predict the US but no US arrives, the mismatch triggers an inhibitory rebound that activates No-US neurons, driving safety learning (Section 2.2.9).

These signals dissociate: (i) a novel situation with an expected outcome (new room, nothing happens) produces novelty without outcome error; (ii) a familiar situation with a surprising outcome (same room, unexpected shock) can produce outcome error with less novelty. In RAM, novelty drives allocation and outcome error drives safety learning. We discuss this in Section 5.

### 2.4 Neural Substrates

Each component of the RAM principle has identifiable neural correlates, which we review here.

#### 2.4.1 Neural reuse in case of familiarity

Considerable evidence suggests that neural ensembles active during encoding are reactivated at retrieval (e.g., Nyberg et al., 2000; Tayler et al., 2012; Tanaka et al., 2014). Importantly, this reactivation is causal for behavior: artificially reactivating encoding-related ensembles can drive context-specific behavioral responses. For example, reactivation of ensembles active during contextual fear conditioning is sufficient to produce a conditioned fear response (freezing) in mice (Liu et al., 2012). This is consistent with our claim that overlap in neural representations drive memory retrieval. As we discuss in Section 5, the stability of such “reuse” appears proportional to encounter frequency: neurons encoding features that are experienced often develop more robust, stable tuning, while those encoding rare features are less reliably reactivated, or reactivated over shorter timescales.

#### 2.4.2 Neural allocation in case of novelty

Two sources can plausibly contribute to the creation of a pool of uncommitted neurons that are repurposed to signal novel representations.

The extreme part of the spectrum is of course neurogenesis, best documented in the dentate gyrus, where young adult-born granule cells preferentially respond to novel stimuli while mature granule cells encode familiar patterns (Nakashiba et al., 2012). However, neurogenesis alone is unlikely to supply sufficient capacity for the volume of novelty an organism processes daily.

A second, and likely more abundant, source of uncommitted neurons is synaptic decay and pruning: memories that are not rehearsed, through recall, replay, or ongoing perceptual reactivation, undergo progressive synaptic weakening. When a neuron’s synaptic configuration has degraded sufficiently, it no longer participates meaningfully in any ensemble and becomes potentially available for re-recruitment through new Hebbian learning. This does not require a return to an embryonic state, but only that prior connections have weakened enough for new patterns to be able to successfully reshape the neuron’s connectivity. A concrete manifestation of this phenomenon is that neurons that lose their original input can be “reassigned”, as well-documented in cases like cortical reorganization after amputation or deafferentation (Merzenich et al., 1984; Pons et al., 1991). Furthermore, this account is consistent with observed representational drift in hippocampal populations (Ziv et al., 2013), where neurons that encode context-specific or temporally specific features show progressive decorrelation over days to weeks.

The picture that emerges is in line with RAM, as reuse and allocation seem inversely related along a single axis: neurons encoding frequently rehearsed features are stably reused and resist reallocation; encoding of rarely rehearsed features decays and these neurons eventually rejoin the uncommitted pool. The same variable— rehearsal frequency—seems to govern both representational stability and neuronal availability.

#### 2.4.3 Episodic uniqueness and timestamping through temporal coding

The reuse-allocation dynamic guarantees that allocation occurs whenever genuinely novel features are present. RAM further assumes that allocation occurs for *every* experience, including repetitions of identical external stimuli. This is plausibly ensured by temporal coding. Time cells in the hippocampus and elsewhere fire in sequences spanning behavioral epochs, providing each moment with a distinct neural signature (MacDonald et al., 2011, 2013). This temporal coding extends to behaviorally relevant timescales: CA1 ensemble patterns decorrelate systematically over hours to days (Mankin et al., 2012), and hippocampal population dynamics timestamp events across weeks, with ensemble similarity decreasing monotonically with temporal distance (Rubin et al., 2015). Human neural recordings confirm that temporal proximity systematically influences pattern overlap, with effects particularly evident in the temporal lobe (Manning et al., 2011).

#### 2.4.4 Episodic event coding

RAM proposes that the brain implements a division of labor between two forms of neural reuse: feature-level neurons reused *across* episodes sharing the same stimulus elements, and event-level neurons allocated for the specific conjunction of features that defines a unique experience. Kolibius et al. (2023) provide direct evidence for this architecture. Recording from single neurons in the human hippocampus, they identified episode-specific neurons (ESNs) that fire selectively during a particular episodic memory, both during encoding and retrieval, coding not for any individual element but for the conjunction of elements composing that episode. Critically, ESNs were dissociable from both concept neurons and time cells. This three-way dissociation mirrors RAM’s architecture: concept neurons correspond to the feature level (reused across events), time cells provide the temporal features that ensure episodic uniqueness, and ESNs correspond to event representations (E_k_) that bind these elements into a sparse episodic index.

Because single or few sparse neurons can reinstate entire associated ensembles through pattern completion (Carrillo-Reid et al., 2016; Robinson et al., 2020), these event neurons serve also as entry points through which a partial cue can reactivate a full memory including its US components, a property central to RAM’s No-US detection mechanism (Section 2.2.9), which we review next.

#### 2.4.5 Biological basis for No-US detection: sparse retrieval and inhibitory rebound

RAM assumes that partial cues reinstate complete event representations through pattern completion: a subset of an event’s neurons, activated by the current cue, can activate the event neuron, which in turn can reactivate its full stored pattern. This property is central to the No-US detection mechanism (Section 2.2.9), where retrieval of a fear event restores the US expectation even in the absence of the US itself. Here we review evidence supporting both elements of this mechanism: sparse pattern completion and an active safety signal when expected outcomes fail to arrive.

That pattern completion from sparse ensembles is plausible is supported by optogenetic evidence: activating hippocampal engram neurons reinstates fear memory recall (Liu et al., 2012), with as few as ∼15 hippocampal neurons sufficient to activate a large ensemble, driving memory-guided spatial behavior (Robinson et al., 2020). In the visual cortex, even a single neuron was shown to reinstate an entire persistent ensemble (Carrillo-Reid et al., 2016).

One possible biological implementation of the US/No-US competition is through mutual inhibition. A top-down US signal, generated by pattern completion but unsupported by external input, cannot sustain itself; the mutually inhibited No-US population rebounds, producing an active safety signal. This would ensure that No-US neurons fire specifically when the situation warrants it: an event that predicts shock has been retrieved, but shock did not come.

Converging evidence supports this account. Tye et al. (2010) showed the existence of amygdala neurons encoding the absence of expected reward. In the aversive domain, Salinas-Hernández et al. (2018) demonstrated that ventral tegmental area (VTA) dopamine neurons fire at the omission of expected shock during fear extinction, and that this signal is causally necessary for extinction learning (optogenetic silencing prevents extinction). This motivates the inclusion of No-US neural pools in our model.

Finally, Herry et al. (2008) identified functionally distinct fear and extinction neuron populations in the basal amygdala exhibiting reciprocal activity patterns during bidirectional transitions between high and low-fear states. These competing populations with inversely correlated activation are consistent with RAM’s mutually inhibited Fear and No-Fear ensembles.

#### 2.4.6 Persistent CS pathway

Our model includes a direct pathway from the CS neurons to the fear/no-fear neurons that bypasses episodic contextual representations. This design choice is motivated by evidence from lesion studies in both rodents and humans. In rats, hippocampal lesions selectively impair contextual fear conditioning while leaving cued (tone) fear conditioning fully intact (Phillips and LeDoux, 1992; Kim and Fanselow, 1992). Romanski and LeDoux (1992) further showed that a direct projection from the auditory thalamus to the lateral amygdala is independently sufficient to support auditory fear conditioning: destroying either the thalamo-amygdala or the thalamo-cortico-amygdala pathway alone had no effect on conditioning, and only combined removal of both routes abolished it. Finally, note that our model’s totally frozen CS pathway is an idealization that models the resistance to change of thalamic fear synapses. Biological persistence is likely graded.

#### 2.4.7 Offline replay and inter-event linking

RAM’s inter-event links form exclusively during offline periods, never during external trials. Hippocampal replay during sleep was first demonstrated by Wilson and McNaughton (1994), who showed that CA1 ensemble firing patterns from waking experience are reactivated during subsequent sleep. Since then, offline reactivation has been observed during both sleep and quiet wakefulness. Furthermore, sharp-wave ripples, brain oscillations during which replay preferentially occurs, were shown to be necessary for memory consolidation during sleep (Girardeau et al., 2009) and for learning during wakefulness (Jadhav et al., 2012).

RAM further predicts that offline reactivation can do more than consolidate individual memories: by co-reactivating ensembles from temporally distant experiences, it can forge new functional links between them. This effect was indeed the key to the success of the Monfils-Schiller effect in our simulations under heavy rehearsal conditions. Zaki et al. (2025) provide direct evidence for this mechanism. Using calcium imaging in hippocampal CA1 of mice, they showed that a strong aversive experience drives offline co-reactivation of not only the recent aversive ensemble but also a neutral-context ensemble encoded two days earlier, and that this co-reactivation is associated with the retrospective spread of fear to the neutral context. Co-reactivation occurred preferentially in quiet wakefulness rather than during sleep.

Finally, Lacagnina et al. (2019) demonstrated that extinction actively suppresses reactivation of fear engram cells during extinction trials while activating a distinct extinction ensemble, supporting RAM’s restriction of inter-event link formation to offline periods rather than online experience.

## 3 Simulation Protocols

### 3.1 Time Steps

All durations are expressed in discrete simulation steps. The model has no intrinsic time unit. Overnight gaps between experimental days are implemented as long silence periods (100 steps), during which weight decay and offline rehearsal operate. Appendix A lists all schedule parameters.

### 3.2 Schedules

We test three families of schedules, incorporating overnight gaps between experimental days, and covering five phenomena: acquisition, extinction, renewal, spontaneous recovery and the Monfils-Schiller effect.

#### Standard schedule

Covers the first three phenomena. Day 1: Acquisition (CS+US in context A). Overnight gap. Day 2: Extinction (CS-alone in context B, *n*_ext_ trials). Overnight gap. Day 3: Tests: Extinction test (CS in B) and Renewal test (CS in A).

#### Spontaneous recovery (SR) schedule

Day 1: Acquisition. Overnight gap. Day 2: Extinction (n_ext_ trials in B). Variable delay (5 or 1500 steps). Test (CS in B). The SR schedule tests whether extinction holds (short delay) or fear returns (long delay), both in the extinction context.

#### Monfils-Schiller schedule

Day 1: Acquisition (CS+US in A). Overnight gap. Day 2: Retrieval (a few CS-only presentations in B). Variable delay (10 or 50 steps). Extinction (*n*_ext_ trials in B). Overnight gap. Day 3: Renewal test (CS in A). Following the original Monfils-Schiller protocol, the retrieval phase occurs in the same context as extinction (B). Since the model operates in discrete time steps, a single animal trial, which unfolds over continuous time, maps to a few discrete presentations (three in our implementation). The schedule tests whether retrieval before extinction prevents fear return at renewal (short delay) or not (long delay).

### 3.3 Evaluation Criteria

At each trial, the model reports Fear (*F*) and No-Fear (*N*) values from outcome competition. The outcome is **Fear** if *F* > N and **No-Fear** if *N* > *F*. The margin |*F* - *N*| measures how decisively the model resolves the competition, with larger margins indicating more robust predictions.

### 3.4 Configuration

Table 2 presents the model configuration used throughout this paper. Key design choices involve: equal weight caps across all pathways (*w*_fear max_ = *w*_no max_ = *w*_cs fear max_ = 1.0), frozen CS pathway after consolidation (*ϕ*_cs min_ = 0), episodic weight decay for non-US events (*w*_decay rate_ = 0.003), and slower decay for US-bearing events (*w*_decay rate us_ = 0 at baseline; robust up to 0.003, Section 4.5).

**Table 2:** Model configuration. Parameters are grouped by function.

| Parameter | Value | Role |
| --- | --- | --- |
| $\beta$ | 1 | Retrieval competition sharpness |
| $\tau_{\text{merge}}$ | 0.80 | Event reuse threshold |
| $k_{\text{time}}$ | 40 | Temporal window width |
| $w_{\text{time}}$ | 0.3 | Temporal weight in overlap |
| $\alpha_{\text{link}}$ | 0.30 | Link learning rate |
| $T_{\text{replay}}$ | 10 | Maximum replay steps per silence period |
| $b_F$ | 0.15 | Fear intrinsic drive |
| $b_N$ | 0.20 | No-Fear intrinsic drive |
| $w_{\text{US} \rightarrow F}$ | 2.5 | Innate US-to-Fear drive |
| $\gamma$ | 1.0 | Mutual inhibition strength |
| $w_{\text{NoUS\_to\_N}}$ | 2.0 | No-US bump magnitude ($\delta_N$) |
| $w_{\text{fear\_max}}$ | 1.0 | Event fear weight cap |
| $w_{\text{no\_max}}$ | 1.0 | Event no-fear weight cap |
| $w_{\text{cs\_fear\_max}}$ | 1.0 | CS pathway weight cap |
| $\phi_{\text{cs\_min}}$ | 0 | CS plasticity after consolidation |
| $T_{\text{virgin}}$ | 20 | Virgin reset window (steps) |
| $w_{\text{decay\_rate}}$ | 0.003 | Per-step decay for non-US events |
| $w_{\text{decay\_rate\_us}}$ | 0.0 | Per-step decay for US events (robust up to 0.003) |
| $n_{\text{ext}}$ | 50 | Number of extinction trials |
| lability_enabled | False | Reconsolidation (disabled; Section 2.2.5) |

## 4 Results

### 4.1 Overview

Table 3 presents the model’s performance on all five target phenomena, each tested at the relevant delay conditions. The model correctly produces the right Fear or No-Fear outcome in every case, with margins ranging from +1.468 to +4.650. We describe the first three below (Figure 3). Spontaneous recovery and the Monfils-Schiller effect will follow.

**Table 3:** Model performance on five fear conditioning phenomena. All test conditions pass with positive margins (min margin +1.468).

| Phenomenon | Expected | $F$ | $N$ | Margin |
| --- | --- | --- | --- | --- |
| Acquisition | Fear | 4.650 | 0.000 | +4.650 |
| Extinction | No-Fear | 0.000 | 3.002 | +3.002 |
| Renewal | Fear | 1.533 | 0.000 | +1.533 |
| SR short (5 steps) | No-Fear | 0.000 | 2.761 | +2.761 |
| SR long (1500 steps) | Fear | 1.468 | 0.000 | +1.468 |
| Monfils-Schiller short (10 steps) | No-Fear | 0.000 | 2.727 | +2.727 |
| Monfils-Schiller long (50 steps) | Fear | 1.533 | 0.000 | +1.533 |

**Figure 3:**
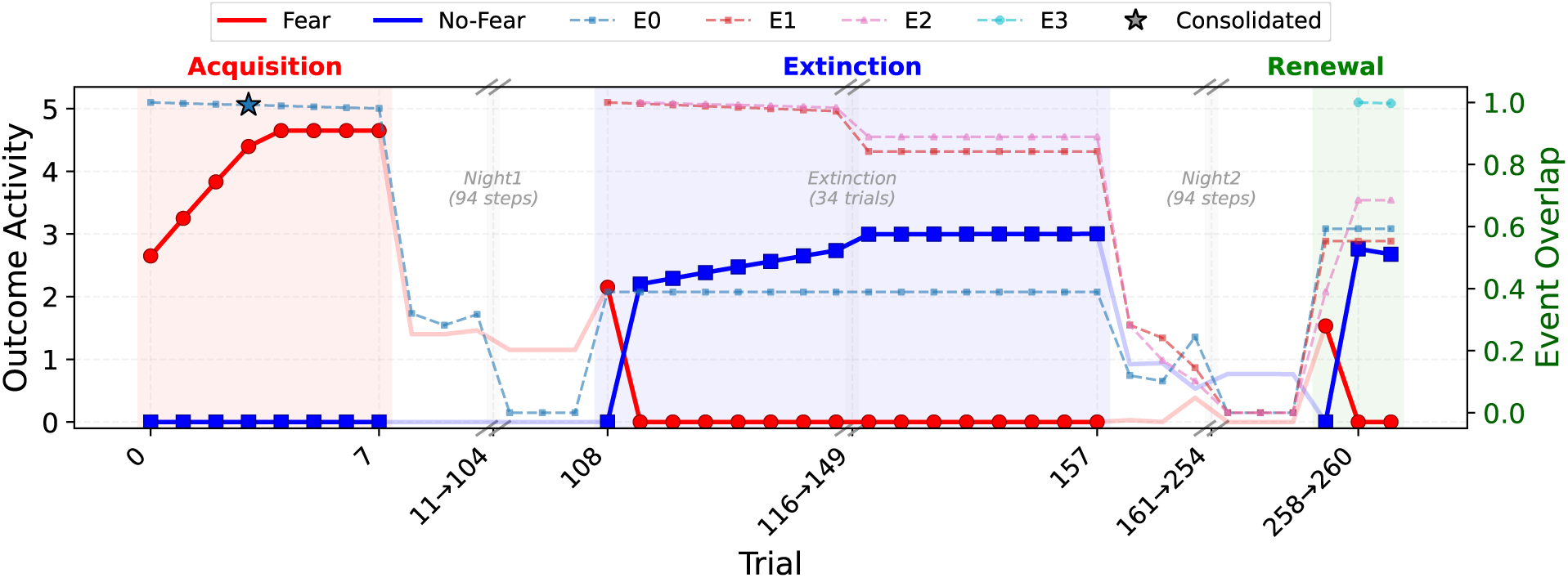
Standard schedule: acquisition, extinction, and renewal. Fear (red) and No-Fear (blue) dynamics with event overlap traces (thin dashed lines). Acquisition drives Fear through *E*_0_; extinction in context B produces No-Fear through new events with No-US markers; renewal in context A restores Fear as *E*_0_ dominates retrieval.

#### Acquisition

A CS-US pairing in context A creates event *E*_0_ (Figure 3) with strong *w*_fear_. The CS pathway also learns fear. At test, *E*_0_ retrieval and the CS pathway drive Fear (margin +4.650).

#### Extinction

Repeated CS-alone presentations in context B allocate new extinction events. The No-US detection mechanism arms after the first trial, and subsequent events incorporate No-US markers. Extinction events accumulate strong *w*_no_ associations. At test in context B, these events dominate retrieval and No-Fear wins (margin +3.002). This demonstrates RAM’s coverage-based allocation: because the extinction cue (context B, no US) has low coverage by *E*_0_ (which was encoded in context A), a new event is created rather than modifying *E*_0_. Safety learning then occurs within the new event through the No-US detection mechanism, leaving *E*_0_’s fear associations intact. These two computations, context-driven event creation and outcome-driven safety learning, are typically collapsed into a single prediction-error signal; committing to RAM separates them (Section 5.1).

#### Renewal

At test in context A, *E*_0_ has full context match while extinction events have weak context A overlap (∼30%). *E*_0_ wins retrieval. Combined with the frozen CS fear pathway, Fear dominates despite prior extinction (margin +1.533; Figure 3). This reproduces the classic finding that extinction is context-dependent (Bouton, 2004).

### 4.2 Spontaneous Recovery

The model reproduces the finding that extinguished fear returns after a long delay but not after a short delay, even when tested in the extinction context (Figure 4).

**Figure 4:**
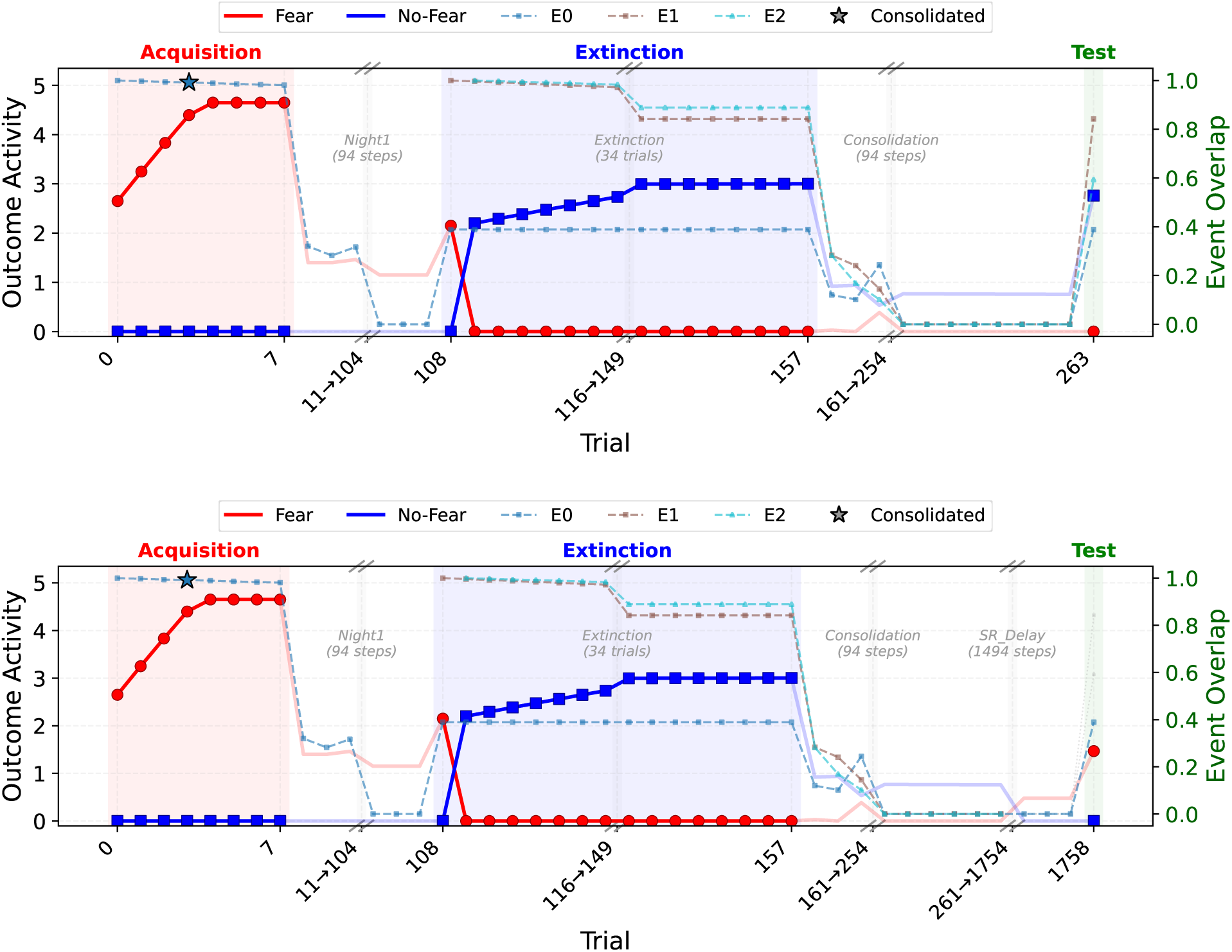
Spontaneous recovery at short and long delays. **Top:** SR short (delay = 5 steps). Extinction events retain nearly full weight; No-Fear wins at test. **Bottom:** SR long (delay = 1500 steps). Extinction events have decayed to ∼1%; the frozen CS pathway drives Fear at test.

**SR short** (delay = 5 steps): Extinction events retain 98.5% of their weights ((1 - *w*_decay rate_)^5^ = (1-0.003)^5^). With 50 trials of accumulated *w*_no_, the combined safety signal overwhelms the frozen CS fear. No-Fear wins with margin +2.761.

**SR long** (delay = 1500 steps): Extinction events retain only 1.1% of their weights ((1 - 0.003)^1500^). The safety signal is nearly eliminated. The frozen CS pathway (*w*_cs fear_ = 1.0) now dominates. Fear wins with margin +1.468 (Figure 4).

At the SR test in context B, three forces compete: (1) extinction events, which have full context B match and whose accumulated w_no_ drives No-Fear; (2) the frozen CS pathway (w_cs fear_ = 1.0), which provides a permanent fear bias independent of event retrieval; and (3) a No-US signal generated when No-US-marked events dominate retrieval. Here, weight decay is the asymmetric lever that affects long delays and after a sufficient number of steps, extinction weights fade to the point where the CS fear bias dominates and fear returns.

### 4.3 Monfils-Schiller Timing Effect

The model can reproduce the finding that retrieval before extinction prevents fear return at renewal, but only when the delay between retrieval and extinction is short (Monfils et al., 2009; Schiller et al., 2010) (Figure 5). Critically, the model predicts that this effect is inherently fragile, consistent with the mixed empirical record (Kredlow et al., 2016). Following the original Monfils-Schiller protocol, the retrieval phase occurs in context B (the extinction context).

**Figure 5:**
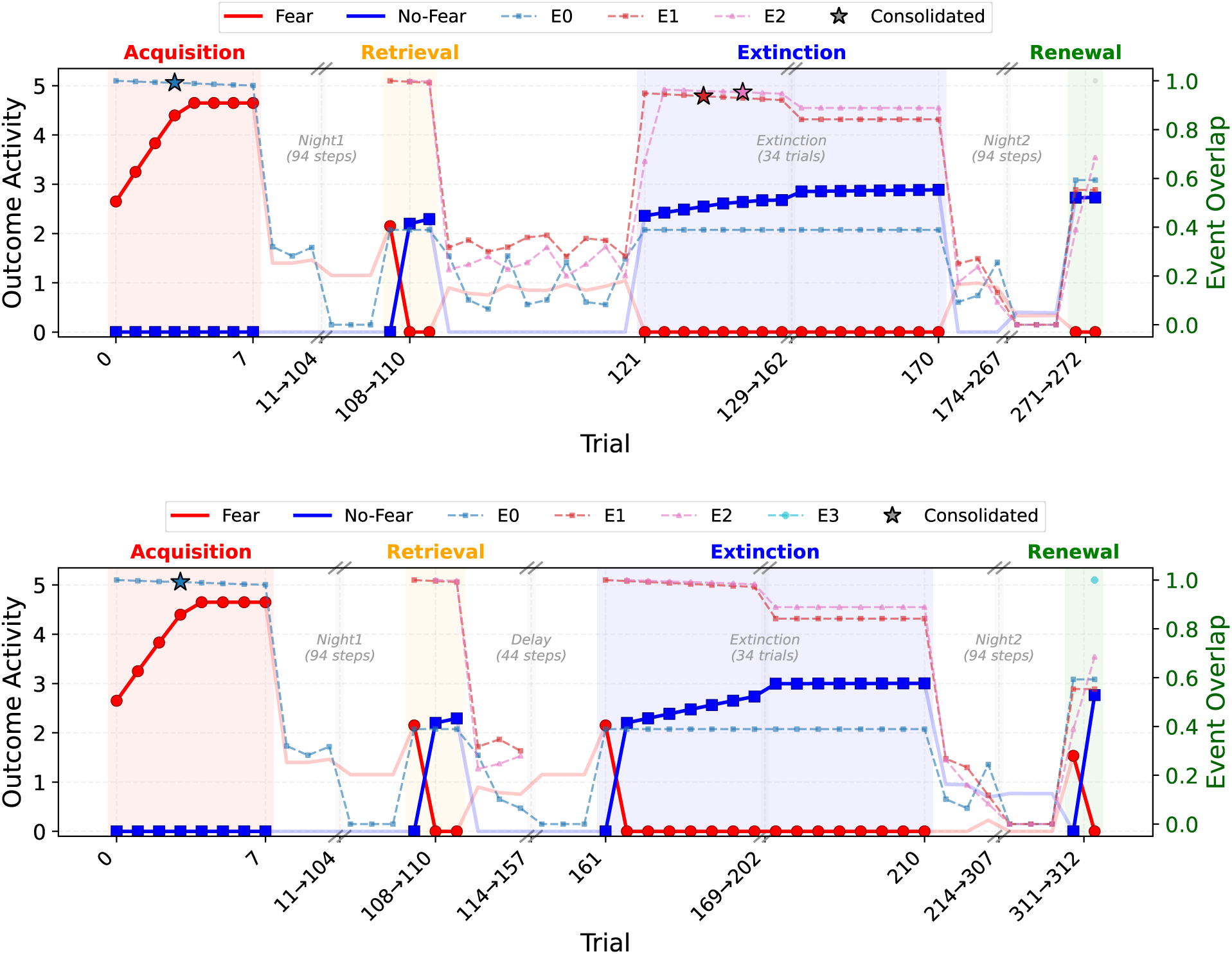
Monfils-Schiller timing effect. Stars (⋆) mark consolidation. **Top:** Short delay (10 steps < *T*_virgin_ = 20). Bridge events *E*_1_ and *E*_2_ survive and consolidate at the first extinction trials, form links during offline replay, and shift renewal toward No-Fear through co-retrieval. **Bottom:** Long delay (50 steps). Bridge events fail to consolidate and are garbage-collected before extinction begins; standard extinction without links, Fear returns at renewal.

**Monfils-Schiller short** (delay = 10 steps < *T*_virgin_ = 20): During the retrieval phase (three CS-only presentations), two events are created: *E*_1_ (a neutral bridge event) and *E*_2_ (which carries No-US features). Both survive the short delay, consolidate at the first extinction trial, and form inter-event links during offline rehearsal. At renewal in context A, *E*_0_ is retrieved by context match, but links amplify *E*_2_ ’s activation. *E*_2_ ’s No-US signal outvotes *E*_0_ ’s US signal, triggering the No-US bump toward No-Fear. The margin for this experiment is +2.727.

**Monfils-Schiller long** (delay = 50 steps ≥ *T*_virgin_ = 20): Both bridge events (*E*_1_, *E*_2_) fail to consolidate within the virgin reset window and are pruned before extinction begins. Without bridges, extinction follows the standard path; new events form that are not linked to *E*_0_. At renewal, *E*_0_ is retrieved alone, and Fear dominates (margin +1.533; Figure 5).

#### Fragility and replay dependence

The Monfils-Schiller effect depends on inter-event links reaching sufficient strength during offline replay to outvote fear at renewal. Setting *α*_link_ = 0 eliminates the effect entirely: without inter-event links, the bridge events created during retrieval cannot be co-retrieved with *E*_0_, and the mechanism fails (the other four phenomena are unaffected; Table 4). Between these extremes, replay duration (*T*_replay_) and per-step inter-event link plasticity (*α*_link_) trade off: we found that weaker per-step plasticity can be compensated by longer replay, and vice versa. This creates a threshold and a prediction tied to total replay intensity: conditions favoring sustained, uninterrupted replay (e.g., quiet isolation) cross the threshold; conditions disrupting replay (e.g., stimulation) do not. This is consistent with the mixed replication record (Kredlow et al., 2016): variability in replay opportunity across laboratories and housing conditions may explain why the effect appears in some studies but not others (Section 5.1).

**Table 4:**
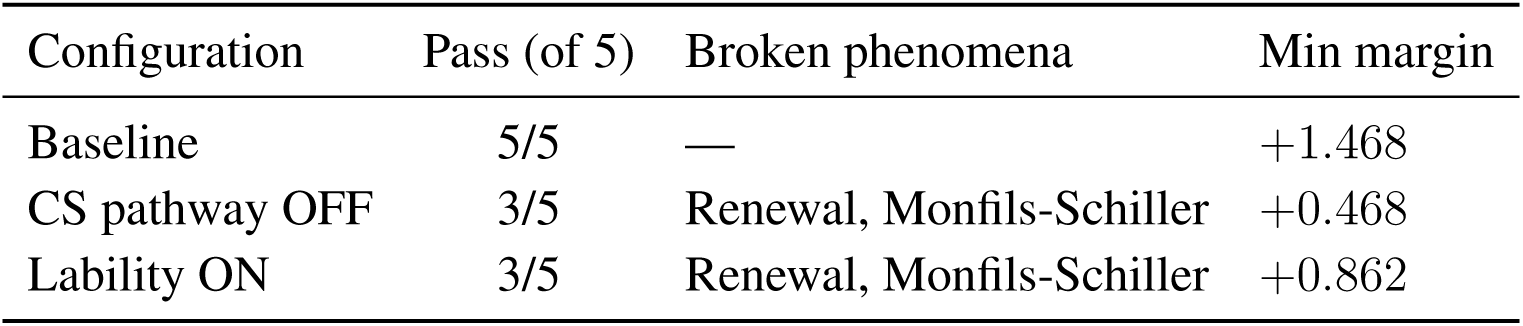
Ablation results. Each row removes or enables one component relative to the baseline.

| Configuration | Pass (of 5) | Broken phenomena | Min margin |
| --- | --- | --- | --- |
| Baseline | 5/5 | — | +1.468 |
| CS pathway OFF | 3/5 | Renewal, Monfils-Schiller | +0.468 |
| Lability ON | 3/5 | Renewal, Monfils-Schiller | +0.862 |

Since the Monfils-Schiller mechanism can operate entirely through appended bridge events and their links, without requiring modification of the original fear memory, this raises a natural question: does reconsolidation of the original trace improve any of the five phenomena?

### 4.4 Pure Append is Most Stable

In the baseline configuration, consolidated events are never modified. Table 4 tests the effect of enabling reconsolidation (lability) and the related effect of disabling the “protective” CS pathway.

Both changes break Renewal and Monfils-Schiller. Disabling the CS pathway alone removes its fear bias directly hence breaking renewal. Lability alone allows consolidated traces (including the CS pathway) to change during extinction hence also weakening renewal (not enough fear memories left).

Since more extinction trials accumulate more safety learning, increasing the pressure on the original fear memory; to disentangle their effects, Figure 6 varies the number of extinction trials across the 2 × 2 CS × Lability factorial (at baseline *β* = 1). The baseline passes all five phenomena from n_ext_ ≥ 30.

**Figure 6:**
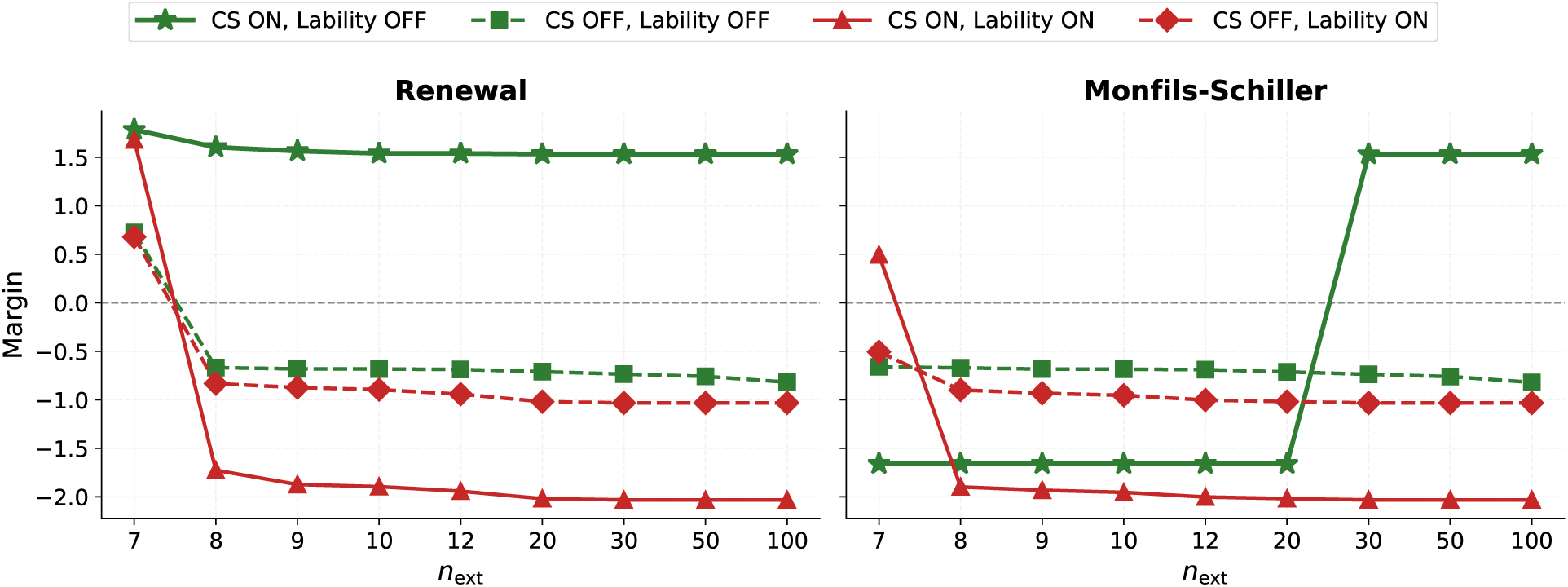
2 × 2 CS pathway × Lability factorial. Each panel shows the margin for one phenomenon as a function of n_ext_. Positive margins indicate the phenomenon is correctly reproduced; negative margins indicate failure. Green: lability OFF; red: lability ON. Solid: CS ON; dashed: CS OFF.

#### The CS pathway protects Renewal against extended extinction

As anticipated, without the CS pathway, *E*_0_ ’s event-level fear alone must sustain Renewal. At *n*_ext_ = 7 this suffices, but any further extinction overwhelms it (*n*_ext_ ≥ 8; Figure 6). This shows that the CS pathway provides a persistent fear bias, independent of event retrieval, that protects both Renewal and Monfils-Schiller long against extended extinction.

#### Reconsolidation restricts without improving

Figure 6 shows that lability ON (even with CS ON) also causes Renewal to fail from *n*_ext_ ≥ 8, the same threshold as CS OFF. Increasing retrieval sharpness (*β*) can rescue the deficit at the baseline (β = 1 and *n*_ext_ = 50): at *β* ≥ 5, all five phenomena pass again (Table 7). However, this introduces a dependence on *β* that does not exist without lability. Without lability, *β* is robust across {1, . . ., 12}; with lability ON, only β ≥ 5 works, and margins are considerably reduced.

Table 8 sheds light on the mechanism. The primary effect is CS pathway neutralization: because CS activation is always 1.0, it crosses the lability threshold at every β, and the CS pathway accumulates w_cs,no_ = 1.0 during extinction. This effectively removes the CS fear bias, placing the model in the same condition as CS OFF analyzed above. Consistent with that analysis, Renewal fails. At *β* ≤ 2, *E*_0_ is additionally modified directly: at β = 1, E_0_’s activation (0.21–0.35) crosses the lability threshold on all 50 extinction trials, accumulating *w*_no_ = 1.0 (our maximum cap), so the original event carries both fear (*w*_fear_) and safety (*w*_no_) associations equally. At β = 2, 24 of 50 trials cross, allowing to to reach *w*_no_ = 0.837. This compounds the deficit but is secondary: at *β* = 3, *E*_0_’s activation drops below the threshold entirely (residual *w*_no_ = 0.068), yet Renewal and Monfils-Schiller still fail because CS pathway neutralization alone suffices. From *β* ≥ 5, *E*_0_ ’s softmax advantage compensates for the lost CS pathway, and all five phenomena pass, paralleling the finding that sharper retrieval partially rescues CS OFF.

We conclude that reconsolidation introduces two restrictions absent from the baseline: a minimum retrieval sharpness (*β* ≥ 5) to compensate for CS pathway neutralization, and a maximum extinction length beyond which Renewal fails (*n*_ext_ ≤ 7; Figure 6). Neither constraint exists without lability. The most stable configuration is hence the extreme form of reuse-and-append: always append new events, never modify existing ones. This almost entirely eliminates the *β* restriction and provides the widest margins.

### 4.5 Parameter Sensitivity

Table 6 summarizes parameter sensitivity. The most constrained parameter is *τ*_merge_, which controls whether a new cue reuses an existing event or creates a new one (working range {0.70–0.85}): below 0.70, extinction cues merge too readily into existing events; above 0.85, the Monfils-Schiller bridge mechanism fails. Weight caps work up to 1.2 (equal across pathways); at 1.6, the model fails. The remaining parameters are robust: *b_N_* (all 6 values pass), *w*_decay rate us_ (all 5 values pass), *w*_time_ (4/5 values pass), and β (1 to 12 without lability; Section 4.4). The protocol parameter *n*_ext_ is robust for the four core phenomena at all tested values; the Monfils-Schiller effect requires *n*_ext_≥ 30. Full results, including two-dimensional interaction grids, are in Appendix C and Appendix D.

## 5 Discussion

Using biologically plausible operations (sparse coding, Hebbian learning, and pattern-matching dynamics), RAM reproduces five key empirical phenomena that motivated the latent cause theory of memory modification (Gershman et al., 2017). Critically, RAM explains these phenomena without needing to modify existing memories and without requiring sharp retrieval competition. The Monfils-Schiller effect, the phenomenon most commonly attributed to reconsolidation, operates through co-retrieval of a bridging event rather than trace modification. This is consistent with evidence that memories can be rescued after reconsolidation blockade (Trent et al., 2015), suggesting the original trace remains intact even when behavioral expression changes.

RAM is broadly consistent with the idea that individual memories are stored separately in the brain, and that retrieval competition is the limiting factor for memory performance (Gershman et al., 2025; Alfei et al., 2026). Retrieval competition arises in part due to gradually drifting neural representations, which affect retrieval by altering neural overlap. Related versions of this idea go back to stimulus sampling theory (Estes, 1955), and play a role in modern memory theories such as the Temporal Context Model (Howard and Kahana, 2002). We have shown how these principles can be brought into alignment (albeit abstractly) with latent cause theory.

### 5.1 Predictions

#### Novelty and prediction error are separate signals

RAM reveals a dissociation within what is computationally modeled as a single prediction-error signal. The model separates two computations: a *novelty signal*, computed by coverage-based allocation, that drives memory formation through virgin neuron recruitment; and an *outcome mismatch signal*, computed by inhibitory rebound when an expected US fails to arrive, that drives safety learning. In standard extinction, these co-occur: the new context provides novelty while US omission provides mismatch, making them difficult to distinguish experimentally. But they are functionally distinct: novelty creates the event (the container), while mismatch provides the safety content (No-US features). The context pre-exposure facilitation effect provides direct evidence for this separation: Fanselow (1990) found that contextual fear conditioning requires sufficient context exposure before shock; with very short placement-to-shock intervals conditioning fails, but prior exposure to the context without shock rescues it. Rudy et al. (2002) demonstrated that this facilitation depends on an intact hippocampal formation, consistent with hippocampal construction of a conjunctive context representation. The pre-exposure phase involves no outcomes and zero prediction error, yet it is essential: the animal must first form the representation (novelty-driven allocation) before it can associate that representation with shock (outcome-driven learning).

#### The Monfils-Schiller effect fails when offline replay is disrupted

The model predicts that the Monfils-Schiller effect is inherently fragile. Simulations show that the bridge network requires unusually strong replay-dependent plasticity, intensive offline rehearsal, or both for the safety signal to outvote fear at renewal (Section 4.3). This is consistent with the difficulty of replicating this effect: Kredlow et al. (2016) found a small, non-significant overall effect in animals (*g* = 0.21, *p* = .30), and Chalkia et al. (2020)’s registered replication found no evidence that reactivation-extinction prevents fear return. Because the model identifies intensive replay as the critical factor, it also explains an interesting potential moderator: Kredlow et al. (2016) found that housing condition explained 30.8% of the variance in effect sizes. Individually housed animals showed a large effect (*g* = 0.78); group-housed animals showed none (*g* = -0.20). Monfils et al. (2009) housed animals individually, placing them in favorable conditions for intensive replay, potentially strengthening inter-event links. Group housing instead introduces competing social stimulation. A direct test would be to house animals individually but with distractions (novel objects, enriched environments) during the post-retrieval delay: the model predicts a weaker Monfils-Schiller effect than in calm, bare housing, dissociating intensive replay from isolation itself.

#### Frequently rehearsed representations drift less

RAM predicts that representational drift rate should inversely correlate with rehearsal frequency across neural populations. Neurons whose representations are frequently reactivated undergo repeated Hebbian strengthening, resisting decay; neurons that are rarely reactivated weaken and become available for re-recruitment. This generates a gradient: feature-selective neurons in early sensory cortex, activated millions of times daily, should be the most stable. Concept neurons (Quiroga et al., 2005), reactivated regularly through daily experience, should drift slowly. Episodic event neurons (Sun et al., 2020), reactivated only during specific recall or replay, should be less stable. Time cells (MacDonald et al., 2011), which encode a particular temporal moment and are unlikely to be specifically rehearsed, should drift fastest. Existing observations are broadly consistent with this ordering: hippocampal ensemble patterns decorrelate over days (Ziv et al., 2013), while sensory cortex representations showed more stability across the tested ranges from weeks to months, especially for simple stimuli (Jeon et al., 2018; Shtoyerman et al., 2000; Marks and Goard, 2021). Face-selective neurons similarly show stability over months (McMahon et al., 2014). More directly, Grosmark et al. (2021) showed that hippocampal place cells recruited to offline reactivation during rest exhibited more stable spatial representations over subsequent days, providing a correlational link between offline reactivation and representational persistence. Even more direct tests could track drift rates across identified cell types (time cells, place cells, concept cells), predicting that drift rate ranks inversely with the frequency at which each representation is naturally reactivated by the animal’s ongoing experience.

### 5.2 Limitations and Future Work

The current model uses abstract feature pools and intersection-based metrics rather than learned representations and true neural dynamics. It assumes an unlimited supply of uncommitted neurons, which contrasts with the finite-brain argument that motivates the principle (Section 2.4). A natural next step is to simulate RAM with realistic neural overlap and competitive allocation in a finite population, testing whether the same principles hold when neurons must be genuinely recycled. Extensions to appetitive conditioning, episodic memory, and other domains remain future work.

## Acknowledgments

SJG was supported by the Kempner Institute for the Study of Natural and Artificial Intelligence, a Polymath Award from Schmidt Sciences, and the Department of Defense MURI program under ARO grant W911NF-23-1-0277.

## Code Availability

Code is available at github.com/bendiogene/reuse and append

## Appendix A Schedule Parameters

Table 5 shows key temporal parameters of our schedules.

**Table 5:** Key schedule parameters in model steps.

| Parameter | Steps |
| --- | --- |
| Overnight gap | 100 |
| Monfils-Schiller short delay | 10 |
| Monfils-Schiller long delay | 50 |
| SR short delay | 5 |
| SR long delay | 1500 |
| Consolidation silence | 100 |

## Appendix B Weighted Overlap and Coverage

The unweighted overlap and coverage formulas in Section 2.2 treat all features equally. In simulations, each feature family *f* ∈ {CS, ctx, time, US, NoUS} carries a weight *w*_f_. Let cue*_f_* and *E_k,f_* denote the neurons in family f that are active in the cue and in event *E_k_*, respectively.

**Weighted overlap** (memory-centric, for retrieval):

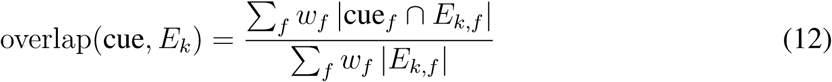

**Weighted coverage** (cue-centric, for allocation):

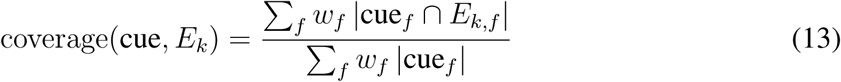

The numerator is the same in both: the weighted count of shared neurons. The denominator differs: overlap normalizes by the event’s weighted size (how much of the stored memory is reactivated), while coverage normalizes by the cue’s weighted size (how much of the current input is explained). All simulations use *w*_CS_ = *w*_ctx_ = *w*_US_ = *w*_NoUS_ = 1.0 and *w*_time_ = 0.3.

## Appendix C Parameter Sensitivity

**Table 6:** Parameter sensitivity summary. Each row shows the values tested, the number passing all five phenomena, and the working range. For *n*_ext_, the four core phenomena (excluding Monfils-Schiller) pass at all tested values; only the Monfils-Schiller effect requires *n*_ext_ ≥ 30.

| Parameter | Tested | Values tested | Pass (of tested) | Working range | Class |
| --- | --- | --- | --- | --- | --- |
| $\beta$ | 10 | {1, 2, 3, 4, 6, 8, 10, 12, 15, 20} | 8/10 | {1, 2, 3, 4, 6, 8, 10, 12} | Robust |
| $n_{\text{ext}}$ | 5 | {12, 20, 30, 50, 100} | 5/5 core; 3/5 all | {30, 50, 100} | Robust (core) |
| $b_N$ | 6 | {0.05, 0.1, 0.15, 0.2, 0.25, 0.3} | 6/6 | {0.05, 0.1, 0.15, 0.2, 0.25, 0.3} | Robust |
| $w_{\text{decay\_us}}$ | 5 | {0, 0.0005, 0.001, 0.002, 0.003} | 5/5 | {0, 0.0005, 0.001, 0.002, 0.003} | Robust |
| cap | 4 | {0.8, 1, 1.2, 1.6} | 3/4 | {0.8, 1, 1.2} | Robust |
| $\tau_{\text{merge}}$ | 8 | {0.5, 0.6, 0.7, 0.75, 0.8, 0.85, 0.9, 0.95} | 4/8 | {0.7, 0.75, 0.8, 0.85} | Narrow |
| $w_{\text{time}}$ | 5 | {0.1, 0.2, 0.3, 0.5, 0.8} | 4/5 | {0.1, 0.2, 0.3, 0.5} | Robust |

Table 7 shows the effect of the retrieval competition parameter β on model performance with and without lability.

**Table 7:** Lability OFF vs ON across β (link alpha = 0.30, n_ext_ = 50). Pass counts are out of five phenomena (and min margins concern only successful phenomena).

| $\beta$ | OFF pass | OFF min margin | ON pass | ON min margin | ON failures |
| --- | --- | --- | --- | --- | --- |
| 1 | 5/5 | +1.468 | 3/5 | +0.862 | Renewal, Monfils-Schiller |
| 2 | 5/5 | +1.456 | 3/5 | +0.851 | Renewal, Monfils-Schiller |
| 3 | 5/5 | +1.446 | 3/5 | +1.446 | Renewal, Monfils-Schiller |
| 4 | 5/5 | +1.439 | 3/5 | +1.439 | Renewal, Monfils-Schiller |
| 6 | 5/5 | +1.431 | 5/5 | +0.213 | — |
| 8 | 5/5 | +1.428 | 5/5 | +0.380 | — |
| 10 | 5/5 | +1.425 | 5/5 | +0.526 | — |
| 12 | 5/5 | +1.423 | 5/5 | +0.633 | — |
| 15 | 4/5 | +1.427 | 5/5 | +0.812 | — |
| 20 | 4/5 | +1.440 | 5/5 | +0.995 | — |

## Appendix D Two-Dimensional Parameter Interactions

Figure 7 shows the margin of the worst-performing phenomenon for each parameter combination. Each cell reports the minimum margin across all five grouped phenomena: a positive value means all five pass (the value is the tightest margin), while a negative value means at least one phenomenon fails (we report the value of the largest failure).

**Figure 7:**
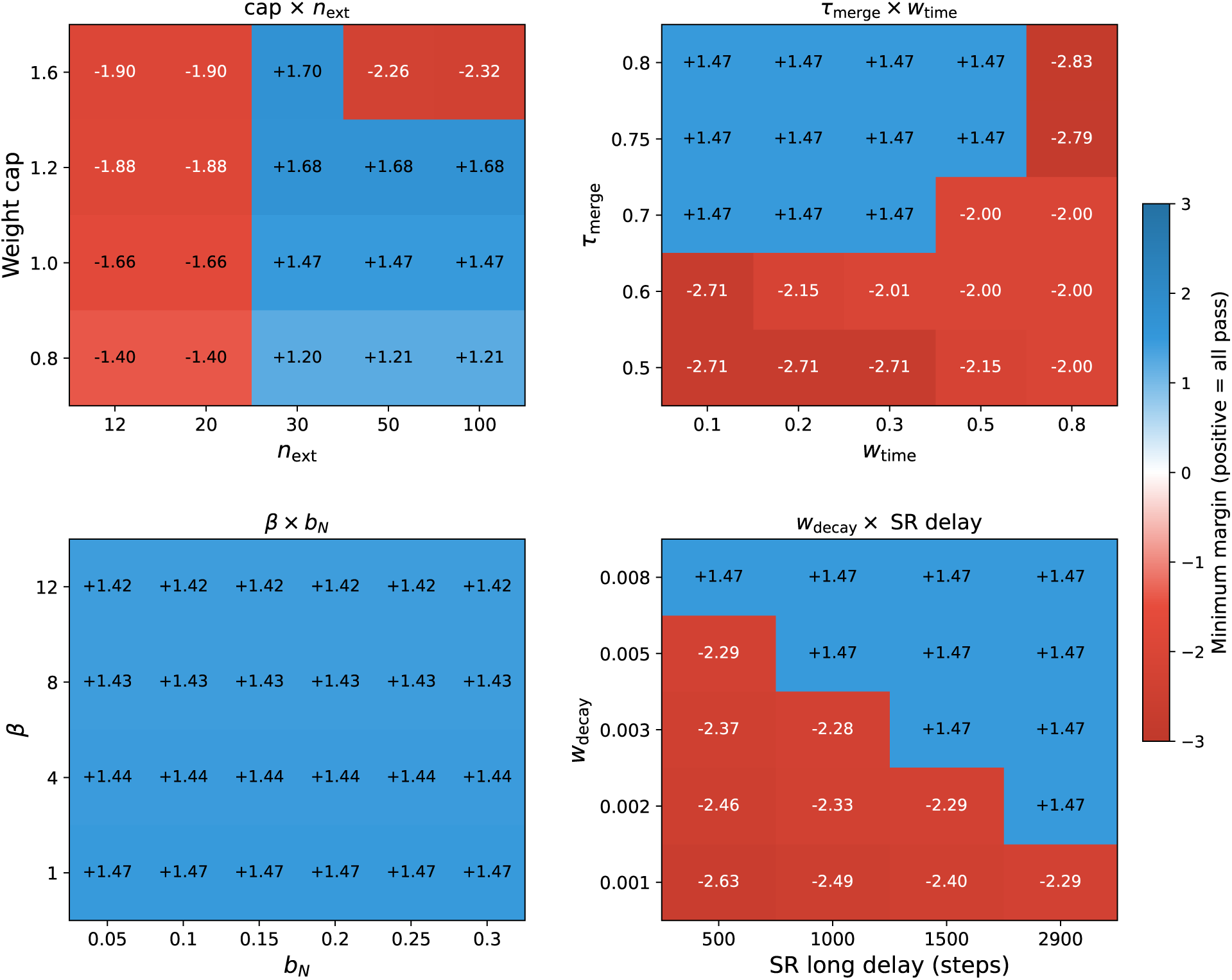
Two-dimensional parameter sweeps. Each cell shows the minimum margin across five grouped phenomena (positive = all pass). The four panels test interactions between weight caps and extinction length, temporal similarity and time weighting, retrieval sharpness and No-Fear intrinsic drive, and weight decay rate and spontaneous recovery delay.

**Cap** × *n_ext_*. For caps of 0.8, 1.0, and 1.2, results are identical: failure at *n*_ext_ ≤ 20 (the Monfils-Schiller effect requires as we saw earlier sufficient extinction trials) and success at *n_ext_* ≥ 30. Cap 1.6 passes only at *n*_ext_ = 30 but fails at *n*_ext_ ≥ 50, likely due to higher caps allowing extinction weights to grow strong enough to suppress fear.

*τ*_merge_ × *w*_time_. The working region is approximately rectangular: *τ*_merge_ ∈ {0.70, 0.75, 0.80} and *w*_time_ ∈ {0.1, 0.2, 0.3}, with *w*_time_ = 0.5 requiring *τ*_merge_≥ 0.75. Both parameters enter the weighted coverage computation that determines whether an incoming cue reuses an existing event or creates a new one, and the two failure modes work in opposite directions.

Below *τ*_merge_ = 0.70, too few events are created during extinction. The first extinction event, allocated before the No-US marker is armed, lacks No-US features. At the baseline threshold (*τ*_merge_ = 0.80), this event is replaced within one trial by a properly formed extinction event carrying No-US features and hence successfully driving safety. At lower *τ*_merge_ thresholds, the firstly created No-US-free event is reused and persists across many next trials: i.e. subsequent extinction cues exceed the lower threshold and reuse it instead of creating a new event. Without No-US features, the residual fear activation from *E*_0_ causes Fear to win the outcome competition on each such trials during extinction, increasing *w*_fear_ through positive feedback until it saturates at the cap. Standard extinction still passes (because a proper safety event (with No US features) eventually forms later), but the Monfils-Schiller effect is the first to fail (at *τ*_merge_ = 0.60) because the brief Monfils procedure cannot overcome the additional fear signal from the contaminated event.

The *w*_time_ failure works in the opposite direction: too many events are created. After overnight gaps (100 steps), temporal overlap between the current cue and any existing event is zero (*k*_time_ = 40 in our simulations), so the temporal term adds to the coverage denominator without contributing to the numerator, deflating coverage. At *w*_time_ = 0.8, this deflation creates 6 events instead of 4, fragmenting the representation and diluting *E*_0_’s activation below the threshold needed to win the Fear/No-Fear competition (Renewal and Monfils-Schiller long fail). At *w*_time_ = 0.5, the deflation is milder but still requires a slightly higher *τ*_merge_ to compensate.

*β* × *b_N_*. Although *β* (retrieval competition) and *b_N_* (tonic No-Fear drive) operate at different stages, they could in principle interact: β determines the activation pattern over events, which feeds into the outcome stage where *b_N_* acts. However, all 24 combinations pass, showing that no such interaction arises within the tested ranges.

*w*_decay_ × **SR delay.** Weight decay rate and SR delay compensate for each other, producing a diagonal boundary as expected. Faster decay requires shorter delays for spontaneous recovery to emerge: at *w*_decay rate_ = 0.008, a delay of 500 steps suffices; at the baseline rate of 0.003, 1500 steps are needed; at 0.002, approximately 2900 steps are required. This follows from the model mechanism: weight decay erodes the extinction no-fear weights over time, and the SR delay determines how much erosion occurs before testing.

Overall, the tested parameter pairs behave as expected intuitively from the model.

## Appendix E E0 Modification Under Reconsolidation

Table 8 shows how the original fear memory *E*_0_ is affected during extinction when reconsolidation (lability) is enabled. Section 4.4 discusses the implications.

**Table 8:**
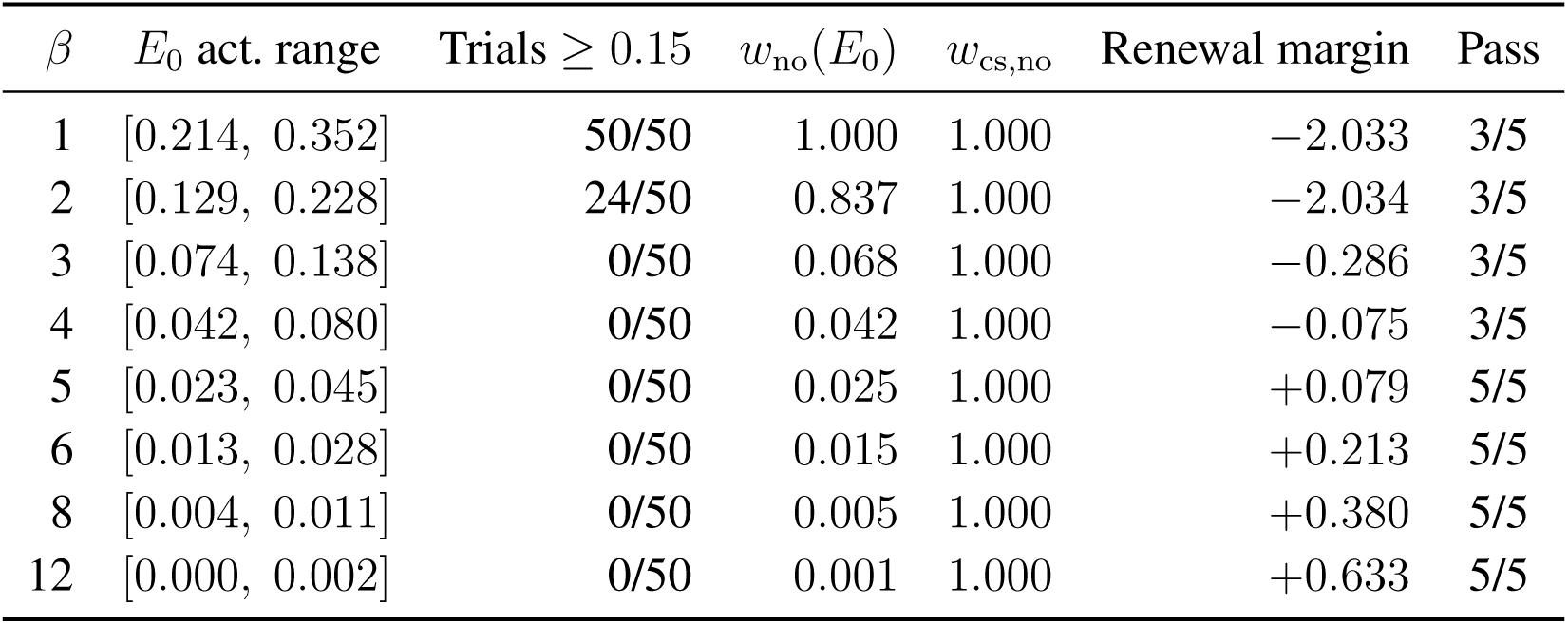
Effect of reconsolidation (lability ON) on the fear memory *E*_0_ and the CS pathway during extinction (*n*_ext_ = 50, lability threshold = 0.15). The CS pathway is fully neutralized at every β (*w*_cs,no_ = 1.0), since cs activation = 1.0 always crosses the lability threshold. With the CS fear bias removed, Renewal depends solely on *E*_0_ ’s event-level fear. At *β* ≤ 4, the softmax is too flat for *E*_0_ to dominate retrieval at Renewal; But starting from *β* ≥ 5, *E*_0_’s activation advantage is sufficient. *E*_0_’s direct modification during extinction (threshold crossing at *β* ≤ 2) is a secondary effect that worsens the Renewal deficit.

